# Integrating Protein and DNA Embeddings for Improving Genome-Wide Transcription Factor Binding Site Prediction

**DOI:** 10.1101/2025.09.15.676319

**Authors:** Shreya Basnet, Jianlin Cheng

## Abstract

Transcription factors (TFs) regulate gene expression by binding to specific DNA sites on genome, making accurate TF binding site prediction critical for understanding gene regulation and downstream phenotypes. Current deep learning methods use only DNA-related information to predict TF binding sites, ignoring the fact that different TF protein sequences and structures recognize distinct DNA patterns. Not leveraging TF information not only limits prediction accuracy but also makes the methods not generalizable to predicting binding sites of new TFs that do not exist in the traning data. Here, we present TransBind, a protein-aware deep learning architecture that integrates DNA sequence information with protein embeddings containing both sequence and structural information derived from a protein language model pretrained on DNA-binding proteins, to improve TF binding site prediction. Through the cross-attention, a TF embedding selectively attends to genomic regions according to its unique binding properties. Evaluated on the data of 690 ChIP-seq experiments spanning 161 TFs across 91 human cell types, TransBind achieves an AUROC of 0.950 and AUPR of 0.371—representing a ≥11.3% relative AUPR improvement over state-of-the-art methods including TBiNet, DanQ, and DeepSEA. The model outperformed existing methods in ≥97.1% of TF–cell type combinations. It also recovered 160 known TF binding motifs in the JASPAR database, providing the biological interpretability of the model. Moreover, the approach enables zero-shot prediction for unseen TFs, demonstrating its potential of generalizing to new, poorly characterized TFs. The source code of TransBind is available at https://github.com/jianlin-cheng/TransBind.

## Introduction

Transcription factors (TFs) are proteins that bind specific DNA sequences to regulate a broad range of cellular processes in almost all species. By activating or repressing target genes, TFs form the backbone of gene regulatory networks, controlling gene expression and protein abundance [7]. For instance, there are over 1,600 predicted TFs in the human genome interacting with millions of potential binding sites [10]. Mapping the transcription factor (TF)–DNA interactome is critical for understanding gene regulation and for linking phenotypic outcomes, such as diseases, to variants in non-coding regulatory regions that disrupt TF binding and alter transcriptional programs [10, 4].

Chromatin immunoprecipitation followed by sequencing (ChIP-seq) [13] is the gold standard for identifying genome-wide TF binding sites, but its scalability is limited by high sequencing costs, substantial material requirements, and dependence on high-quality TF-specific antibodies. These constraints have driven the development of computational approaches to predict TF-DNA binding directly from genomic sequence.

Early models, including position weight matrices [18], hidden Markov models [12, 11], and support vector machines [23, 6] could capture DNA sequence preferences of TFs but had a high false-positive rate and relied heavily on hand-crafted features. Deep learning methods transformed the field by enabling end-to-end learning from raw DNA sequences. Convolutional neural networks (CNNs), such as DeepBind [20], learned sequence motifs directly, while multi-task models such as DeepSEA [22] jointly predicted hundreds of chromatin features, improving performance through shared representations. Hybrid CNN–RNN architectures like DanQ [16] captured long-range dependencies, and attention-based methods such as TBiNet [15] and DeepGRN [5] improved interpretability by focusing on the most informative sequence regions.

Despite these advances, almost all existing approaches use only DNA sequences and/or other genomics data as input without using TF information at all, overlooking critical determinants of binding specificity such as TF’s amino acid sequence and three-dimensional structure [14, 19]. This omission limits both their predictive accuracy and generalization, particularly for TFs with limited binding data. It makes the existing method lack the capability of predicting binding sites for new TFs never seen before (i.e., zero-shot prediction) because they cannot leverage the sequence and structural properties of TFs and the similarity between TFs to generalize. Therefore, it is important to integrate DNA and TF information to improve the prediction of TF binding sites on genome.

Protein language models (PLMs) offer a promising way to represent TF information. Models such as ESM-2 [17] learn rich, contextualized representations from millions of protein sequences, capturing structural and functional properties. Domain-adapted variants, such as ESM-DBP [21], specialize in DNA-binding proteins, embedding biochemical features directly relevant to TF–DNA recognition. However, there is no prior work to integrate the protein-derived embeddings with DNA sequence features to enable dynamic TF-DNA interaction modeling and robust generalization to unseen TFs.

Here, we present **TransBind**, a protein-aware deep learning architecture that integrates DNA sequence encodings with TF protein embeddings via a cross-attention mechanism. The design allows each TF to selectively attend to genomic regions based on its unique binding properties. Evaluated on a large dataset, it significantly outperforms the state-of-the-art deep learning methods. Moreover, our approach supports zero-shot prediction for TFs unseen during training. By uniting protein and DNA features, the approach advances both the predictive performance and biological interpretability in TF-DNA binding prediction.

## Materials and Methods

### DNA Data

To train and evaluate transcription factor (TF)–DNA binding prediction, we used EPBDXDNA dataset [8], derived from ENCODE (Encyclopedia of DNA Elements) ChIP-seq experiments. EPBDXDNA includes 690 TF-cell type experiments spanning 161 TFs across 91 human cell types. Multiple experiments per TF in different cell types capture diverse cellular contexts and experimental conditions. The EPBDXDNA dataset is the updated version of DeepSEA dataset [22] based on the recent genome assembly (hg19). The TF-binding peaks were called by the standardized processing pipeline provided by the ENCODE Analysis Working Group (AWG).

Based on the GRCh37 (hg19) reference genome [9], we partitioned the genome into non-overlapping 200bp bins. A bin was labeled positive if at least 50% of its length overlapped a ChIP-seq peak [22]; otherwise, it was labeled negative. To provide a sufficient context for TF-binding site prediction, each bin was extended by 400bp on either side, resulting in a 1,000 bp input DNA sequence.

In cases where multiple peaks from the same experiment overlapped a single bin, we resolved duplicates by retaining the first instance and merging their labels. After de-duplication, we obtained 1,903,712 unique labeled bins. Removing sequences containing ambiguous nucleotides (‘N’) yielded a final set of 1,903,668 bins, covering approximately 12.6% of the genome. Reverse complement sequences of the bins were also incorporated into the training, validation, and test sets, effectively doubling the total number of samples to 3,807,336.

Each 1,000 bp genomic sequence bin was one-hot encoded as a 1,000 × 4 matrix representing the nucleotides A, C, G, and T. For each of the 690 TF-cell type experiments, a binary value was assigned to each bin indicating whether it has a TF binding site in the experiment or not (positive or negative). This resulted in a binary label vector of length 690 for each bin, indicating in which TF-cell type experiments the bin is positive or negative.

To ensure consistency with prior work, we adopted the chromosome-based data split used in DeepSEA [22]. The bins of Chromosomes 8 and 9 were held out for testing, the bins of chromosome 7 were used for validation, and the bins of the remaining autosomes plus chromosome X (excluding Y) were used for training. The resulting data distribution is summarized in **Table 1**.

**Table 1.** Chromosome-based data partitioning used for training, validation, and testing.

| Split | Chromosome(s) | Bins | % |
| --- | --- | --- | --- |
| Train | 1–22, X (excl. 7–9) | 3,272,910 | 85.96 |
| Validation | 7 | 203,784 | 5.35 |
| Test | 8, 9 | 330,642 | 8.69 |
| **Total** |  | **3,807,336** | **100.00** |

**Figure 1.** illustrates the distribution of TF binding frequencies across the 690 ChIP-seq TF-cell type experiments in the training, validation, and test sets, respectively. The cumulative distribution graph demonstrates severe class imbalance in TF binding frequencies: in approximately 50% of the TF-cell type experiments the TF binds to fewer than 1% of bins, while in 75% it binds to fewer than 2% of bins. Only a small fraction of TFs exhibit binding activity above 3% frequency. This pronounced skew toward low binding frequencies is consistent across all three dataset splits (train, validation, and test), confirming that positive binding sites are very rare. With positive examples comprising only 1.44% bins across 690 TF-cell type experiments, the dataset is therefore highly imbalanced, with the vast majority of examples being negative.

**Fig. 1.**
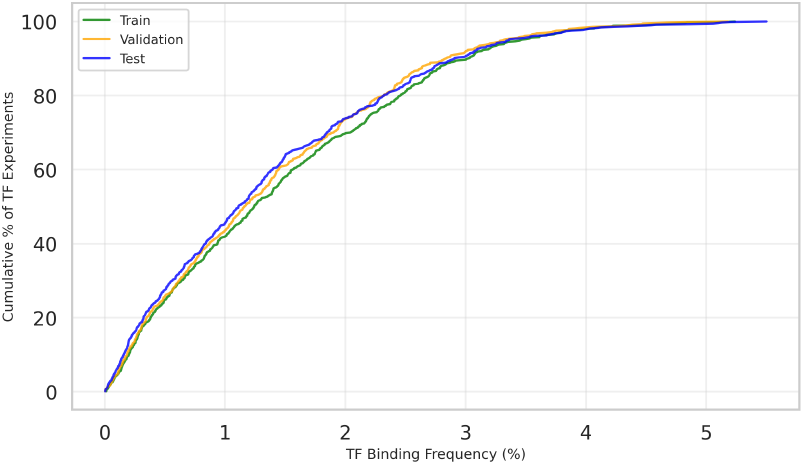
Cumulative distribution of TF binding frequencies in the training, validation, and test sets, respectively. The plot shows the percentage of the TF-cell type experiments (y-axis) with binding frequencies at or below each threshold (x-axis), demonstrating that most TFs bind to a very low proportion of genomic regions while only a small fraction of TFs exhibit substantial binding activity (> 3%) in each set.

### Transcription Factor Data

The amino acid sequences of the 161 TFs were retrieved from UniProt. We used ESM-DBP, a domain-adapted protein language model trained specifically for DNA-binding proteins [21] to generate the embedding for each TF as features. ESM-DBP extends the general-purpose protein language model ESM2 [17] by fine-tuning the pretrained ESM2 on 170,264 non-redundant DNA-binding protein sequences from UniProtKB [1], thereby incorporating TF-specific knowledge. Each TF sequence was processed with ESM-DBP to produce an embedding of shape *L* × *d*_2_, where *L* is the sequence length and *d*_2_ is the embedding dimension. For TF-DNA binding site prediction, we derived a single *d*_2_-dimensional feature vector for a TF by averaging the embedding across the sequence length.

### Deep Learning Model for TF-DNA Binding Site Classification

All the existing deep learning were trained to predict binding sites of a predetermined set of TFs (or TF-cell type combinations) with the availability of some experimentally determined DNA binding sites that can be used for training. The problem is often formulated as a multi-label classification problem, i.e., predicting if a DNA bin contains a binding site of each TF in the set. It is treated as multi-label classification because one bin may be the binding site of multiple TFs. Following this paradigm, we designed the architecture of TransBind to integrate protein and DNA information to predict the binding sites of a set of TF experiments (i.e., 690 TF-cell type experiments in this study) with labeled training data (**Figure 2**). It has three modules: (1) DNA Sequence Encoder, (2) Protein Encoder, and (3) Bimodal Feature Aggregation for TF-DNA Binding Prediction, which are described in detail below.

**Fig. 2.**
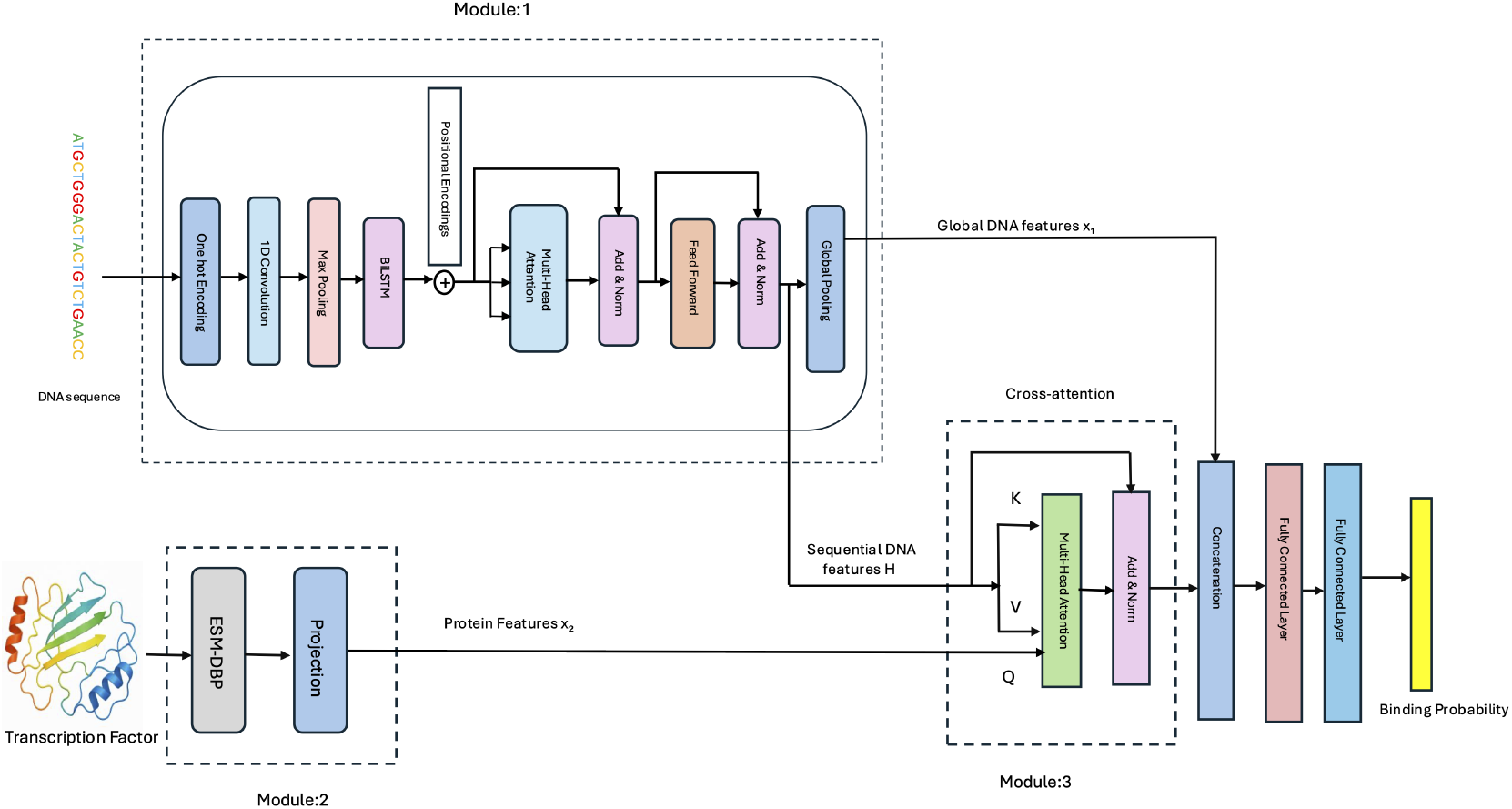
Overview of the TransBind architecture. Module 1 encodes the DNA sequence of a bin using convolutional, BiLSTM, and Transformer layers to generate a contextual DNA representation (embedding). Module 2 encodes TF sequences using pretrained ESM-DBP embeddings. Module 3 applies the cross-attention between TF and DNA features to predict TF-DNA binding probability.

#### Module 1: DNA Sequence Encoder

The binary matrix of shape 1000 × 4 encoding the DNA sequence of each bin is first transposed to the shape 4 × 1000 to match the input format of the following convolutional layer (see Module 1 in **Figure 2**). The transposed matrix is passed through a one-dimensional convolutional layer with 320 filters (kernels) and a kernel size of 26, designed to capture local sequence motif patterns. The convolutional output is followed by a ReLU activation function and a one-dimensional max-pooling operation with a window size of 13, reducing the length of generated hidden features while retaining the strongest activations.These operations progressively reduce the sequence length from the original 1000bp input to an effective sequence length of L=75 positions.

The pooled features are then entered into a two-layer bidirectional long- and short-term memory block (BiLSTM) with a hidden layer of 160 hidden nodes in each direction, resulting in a *d*_1_-dimensional contextualized representation at each sequence position (*d*_1_ = 320)). This enables the model to capture long-range dependencies between features from both upstream and downstream sequence contexts.

The sinusoidal positional encodings are added to the BiLSTM output to be used as input for the following transformer encoder.The transformer encoder has a multi-head self-attention layer with 16 attention heads and a feed-forward layer with 1,024 hidden nodes, allowing the model to attend informative features globally.

Finally, the DNA encoder produces two representations: 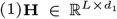, the position-wise features from the transformer encoder (with *L* = 75 and *d*_1_ = 320), which preserve spatial information along the sequence; and 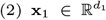, a global embedding obtained by applying average pooling to **H**, serving as a compact summary of the sequence.

**H** is integrated with the protein embeddings generated in Module 2 through the cross-attention in Module 3, while **x**_1_ is concatenated with the output of the cross-attention to predict TF-DNA binding in Module 3.

#### Module 2: Protein Encoder

TransBind leverages ESM-DBP, a domain-adapted variant of ESM-2 fine-tuned for DNA-binding proteins, to generate the embedding for each TF, which can capture sequence and structural features relevant to DNA–protein interactions (see Module 2 in **Figure 2**). For each of the 161 unique TFs in the dataset, a fixed-length embedding of dimension *d*_2_ (*d*_2_=1280) is generated by ESM-DBP. This fixed-length embedding is achieved by mean pooling over the per-residue embeddings generated by ESM-DBP.

To facilitate the integration with the DNA sequence embeddings from Module 1, each protein embedding is projected into a *d*_1_-dimensional latent space using a learnable linear transformation:

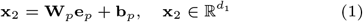

where 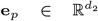 is the original ESM-DBP protein embedding, 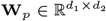 is a trainable projection matrix, and 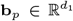 is a bias term. The resulting protein vector **x**_2_ has the same dimensionality of the DNA embedding to facilitate the subsequent cross-modal interaction in Module 3.

#### Module 3: Bimodal Feature Aggregation for TF-DNA Binding Prediction

A key innovation of TransBind lies in a cross-attention mechanism that enables bimodal interaction between DNA and TF embeddings (features) (see Module 3 in **Figure 2**). Biologically, we want each TF to “scan” the DNA sequence and focus on relevant binding regions. We implement this using cross-attention where the TF embedding acts as the query, while the contextualized DNA embeddings serve as both keys and values.

Formally, let 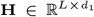 denote the contextualized DNA embeddings from Module 1, where *L* is the effective sequence length after convolution and pooling, and let 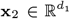 be the projected TF embedding from Module 2. The cross-attention is computed as:

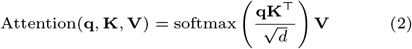

where **q** = **x**_2_**W**_*q*_ ∈ ℝ^1×*d*^ is the TF query vector, and **K** = **HW**_*k*_ ∈ ℝ^*L*×*d*^ and **V** = **HW**_*v*_ ∈ ℝ^*L*×*d*^ are the key and value matrices derived from the DNA embeddings. Here, **W**_*q*_, **W**_*k*_, 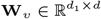 are learnable projection matrices, and *d* is the attention dimensionality.

The output of the attention mechanism is a TF-aware DNA summary vector **z** ∈ ℝ^*d*^, capturing binding-relevant information from the DNA sequence conditioned on the TF. To enrich the representation, **z** is concatenated with the global DNA embedding **x**_1_ (from Module 1), and the resulting vector is passed through two fully connected layers with a ReLU activation on the first layer, followed by a sigmoid function to convert raw logits into binding probabilities between 0 and 1 for the final multi-label prediction.

### Training and Evaluation

TransBind was implemented using PyTorch Lightning. It was trained on the training data and its hyperparameter optimization was performed on the validation data using Optuna (see Supplementary Note S1 for details). The final optimized model employed a 1D CNN (320 channels, kernel size 26), 2-layer bidirectional LSTM (160 hidden units), and transformer encoder with 16 attention heads. The cross-attention mechanism also has 16 heads. Training utilized binary cross-entropy with logits loss, the AdamW optimizer (learning rate 3.28 × 10^−4^, weight decay 0.028) with cosine annealing scheduling over 60 epochs, dropout regularization (*p* = 0.088), and batch size 256.

The performance of TransBind and other baseline methods was blindly evaluated using the Area Under the Receiver Operating Characteristic curve (AUROC) and the Area Under the Precision and Recall Curve (AUPR) metrics across all 690 TF-cell type experiments in the test data.

### Deep Learning Model for Zero-Shot TF-DNA Binding Site Prediction

To enable prediction of DNA binding sites for TFs unseen during training, we reformulate TF–DNA binding prediction from a multi-label classification problem in the previous section into a general binary classification task. Instead of simultaneously predicting binding probabilities for a preselected set of TFs given a DNA bin, the new model addresses a fundamental question: given an arbitrary single TF and a DNA bin, will they bind? Except for the difference in the final output layer in Module 3, the new model (TransBind zeroshot) is the same as TransBind for multi-label classification. Rather than learning binding patterns of a predetermined set of TFs, TransBind zeroshot is trained to capture the universal principles of any protein–DNA interactions that underlie binding affinity and specificity.

One key architectural insight is that the cross-attention mechanism in TransBind zeroshot operates as a shared module for processing individual TF–DNA pairs. During training, each DNA sequence is paired with multiple TF embeddings and vice versa, enabling the model to learn sequence motifs and structural determinants of compatibility. This setup compels the model to extract generalizable features of protein–DNA recognition—such as amino acid–nucleotide contact preferences, structural complementarity, and physicochemical compatibility.

To further promote generalization, we applied regularization strategies in training: randomly masking entire TF embeddings with probability 0.05 and injecting small Gaussian noise (*σ* = 0.01) into protein representations. These techniques discourage reliance on specific TF identities and improve robustness to variability in protein features.

Consequently, once trained, the model can predict binding for entirely novel TFs by simply providing their ESM-DBP embedding together with a DNA sequence, without requiring additional training or fine-tuning. This zero-shot prediction capability substantially improves the model’s flexibility and applicability, offering a general framework for TF–DNA binding prediction across arbitrary TFs.

For training and validation, we used the same dataset as TransBind. The test of zero-shot prediction was blindly carried out on the data of three transcription factors (HNF1A, ATF4, and FOXA3) excluded from training and validation. This setup ensured the model was tested on truly unseen TFs, directly assessing its ability to generalize to novel TF–DNA interactions.

## Results

### Comparison with State-of-the-art TF-DNA Binding Prediction Methods

To rigorously evaluate TransBind, we adopted the same benchmarking framework used in previous state-of-the-art methods, including DeepSEA [22], DanQ [16], and TBiNet [15], to compare them on the same test dataset. For fairness, we retrained all baseline models (DeepSEA, DanQ, TBiNet, and DNABERT-2) on our training dataset. We report their area under the receiver operating characteristic curve (AUROC) and area under the precision-recall curve (AUPR). AUROC measures a model’s ability to distinguish between positive and negative classes across varying thresholds, while AUPR emphasizes performance (precision and recall) on the positive class, making it particularly informative for highly imbalanced datasets such as TF-DNA binding data.

As shown in **Table 2**, TransBind outperforms all baselines across both metrics, achieving an AUROC of 0.950 and an AUPR of 0.371. Compared to the second most accurate predictor - TBiNet, this represents an absolute improvement of +0.012 in AUROC and +0.0376 in AUPR, corresponding to a relative gain of over 11% in AUPR. Moreover, we conducted paired t-tests comparing TransBind against TBiNet across all 690 transcription factor-cell type combinations. The results demonstrate highly significant improvements over TBiNet: AUROC (p=2.48 × 10^−91^, Cohen’s d=0.90) and AUPR (p=2.09 × 10^−60^, Cohen’s d=0.69).

**Table 2.** AUROC and AUPR of TransBind and other methods on the test dataset.

| Method | AUROC | AUPR |
| --- | --- | --- |
| DeepSEA | 0.893 | 0.250 |
| DanQ | 0.925 | 0.306 |
| TBiNet | 0.940 | 0.335 |
| finetuned DNABERT-2 | 0.918 | 0.296 |
| **TransBind** | **0.950** | **0.371** |

**Figure 3** presents the average ROC and PR curves across all 690 TF-cell type combinations across all methods. The ROC curves show TransBind’s ability to achieve higher true positive rates while maintaining lower false positive rates compared to baseline methods. The PRC curves indicate TransBind obtains higher precision and recall across board.

**Fig. 3.**
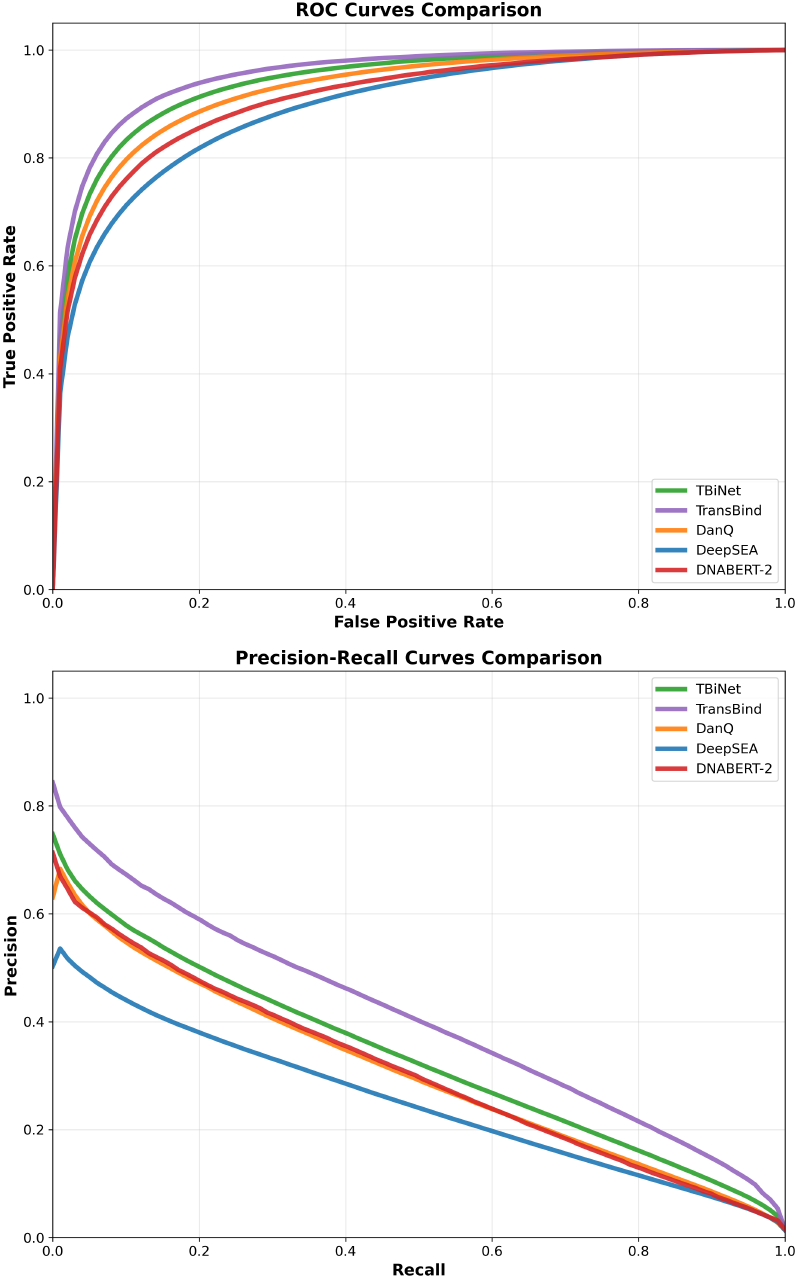
Average ROC curves (top) and Precision-Recall curves (bottom) across all TF-cell type combinations for all methods. TransBind demonstrates superior performance in both metrics, with particularly notable improvements in the precision-recall space.

To assess the performance across individual TFs, we conducted a fine-grained comparison between TransBind and TBiNet across all 690 TF–cell type combinations in the test dataset (**Figure 4**). TransBind outperformed TBiNet in 96.1% (663/690) of cases in terms of AUROC and 97.1% (670/690) in terms of AUPR, underscoring its robustness and broad applicability. The individual performance curves for each TF-cell type experiment are provided in **Supplementary Figures S1 and S2**.

**Fig. 4.**
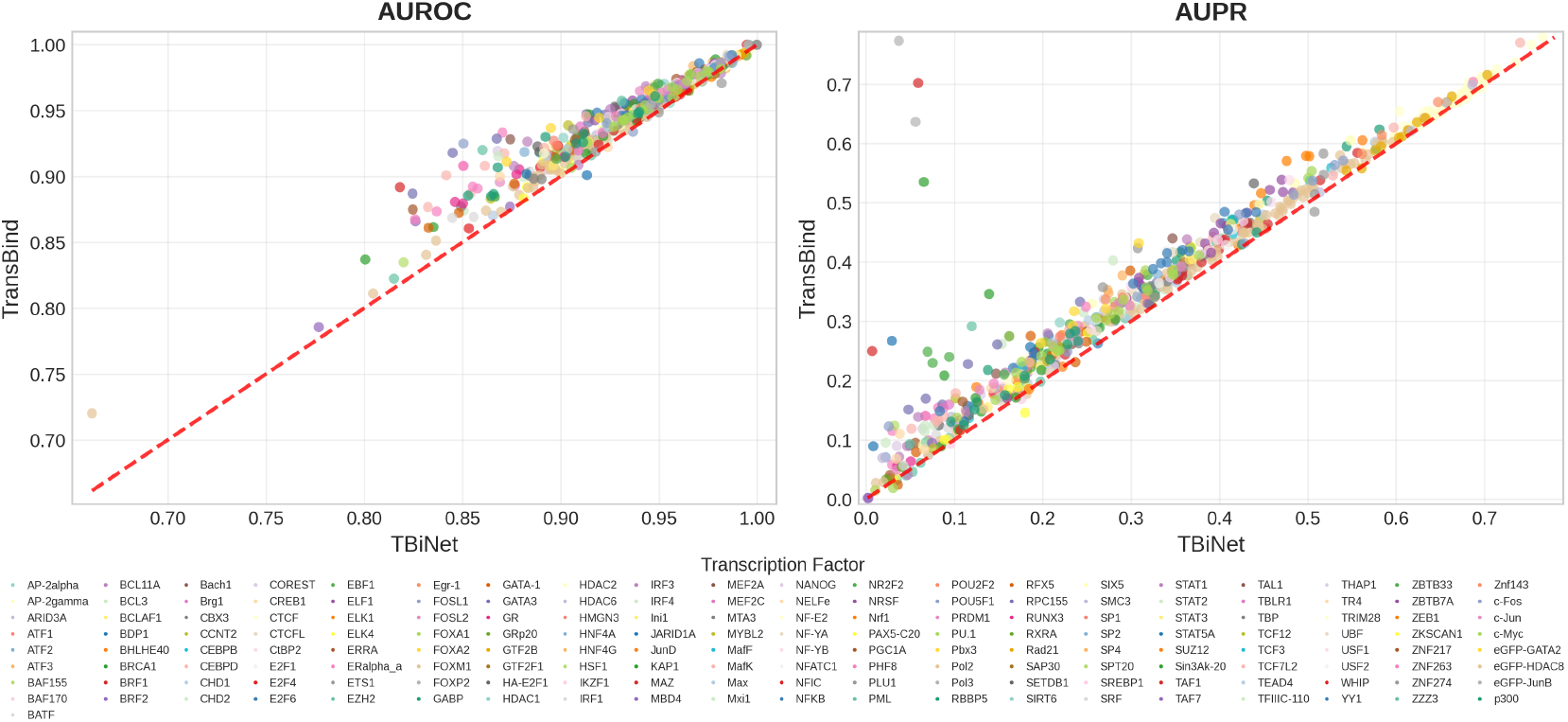
Performance comparison of TransBind vs. TBiNet across 690 TF-cell type combinations involving 160 TFs and 91 cell types in the test set. Each point represents a TF-cell type experiment in terms of AUROC (left) and AUPR (right). Points above the diagonal line indicate better performance by TransBind. Each color denotes each unique TF.

The improvement is especially meaningful in biological contexts where the underlying data is highly imbalanced and the cost of false positives is high. For example, in many challenging cases where TBiNet failed, TransBind achieved dramatic gains, such as Pol3 in K562 (AUPR 0.037 → 0.773) and BRF1 in K562 (0.059 → 0.702). Even in many easier cases where the baseline performance was already good, TransBind still yielded notable improvements, e.g., CTCF in H1-hESC (0.604 → 0.654) and ZNF274 in HepG2 (0.518 → 0.583). Higher AUPR values translate into more precise identification of true binding sites, enabling more efficient downstream experimental validation.

### Biological Interpretation of TransBind

To evaluate whether TransBind captures genuine regulatory mechanisms that are biologically explainable, we systematically analyzed the learned convolutional filters with respect to TF binding motifs. We extracted the trained weight matrices of all 320 kernnels in the first CNN layer in TransBind and queried them against the known TF binding motifs in the JASPAR [3] database using TOMTOM [2] similarity analysis.

TransBind achieved excellent motif discovery performance, learning 160 biologically meaningful kernels matching known TF binding sites in JASPAR with statistical significance (*p <* 0.05). Twenty-seven kernels reached highly significant matches (*p <* 10^−6^), with the strongest discovery achieving *p* = 1.29 × 10^−8^ for the glucocorticoid response element binding protein Gmeb1.

The discovered motifs encompass diverse regulatory pathways essential for cellular function. The most prevalent binding motifs belong to CTCF (36 kernels), the master organizer of chromatin architecture and genomic looping. Additional discoveries span critical regulatory programs, such as developmental patterning (Six4, Lhx1), immune response (Nfatc2), metabolic control (HNF1A), and chromatin remodeling factors. The breadth of the discovery demonstrates that TransBind learned fundamental principles of gene regulation across multiple biological processes.

**Figure 5** presents sequence logo comparisons between TransBind’s top motif discoveries and their JASPAR references. The striking visual correspondence—particularly for Six4, CTCF, and Nfatc2—provides compelling evidence that TransBind captured authentic regulatory grammar rather than dataset-specific patterns. Importantly, these motifs were learned de novo from sequence data alone, without explicit knowledge of binding site locations. Interestingly, the motifs were reflected in the weight matrices of the kernels in the first hidden layer of TransBind, even though the kernels were not explicitly designed to capture the motifs.

**Fig. 5.**
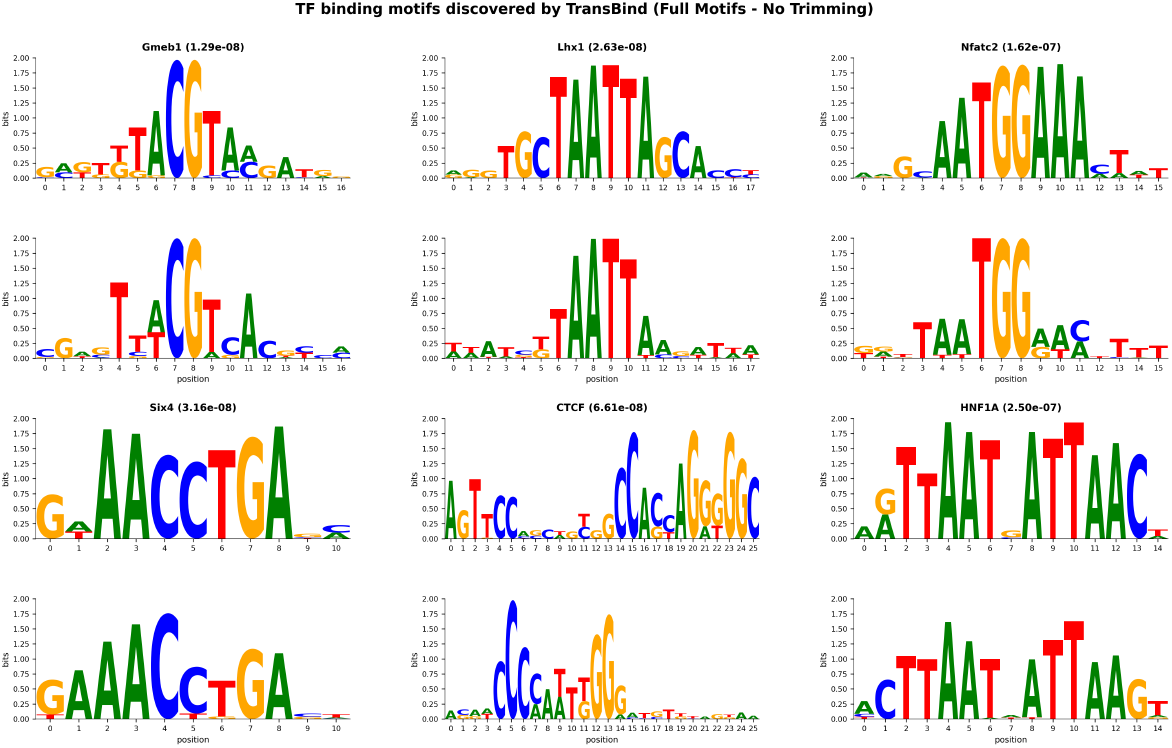
Transcription factor binding motifs discovered by TransBind. Each pair of logos for a TF shows the JASPAR reference motif (top) compared with the corresponding motif learned by TransBind (bottom). TF names and statistical significance (p-values) are shown above each pair. All discoveries have *p <* 2 *×* 10^−7^, demonstrating that TransBind successfully learned biologically meaningful binding patterns for diverse transcription factor families including chromatin organizers (CTCF), homeobox factors (Six4, Lhx1), immune regulators (Nfatc2), and metabolic factors (HNF1A, Gmeb1).

The results establish that TransBind’s architecture not only achieves state-of-the-art prediction accuracy but also provides unprecedented biological interpretability in the regulatory genomics domain.

### Zero-shot TF-DNA Binding Prediction for Unseen Transcription Factors

Existing deep learning methods can only predict binding sites for one or a set of preselected TFs that are included in the training data because they only use DNA information as input to make predictions. In contrast, our approach (e.g., TransBind zeroshot) has the potential to generalize to predict binding sites for new TFs never seen before because it takes both TF and DNA sequence information as input to assess their match. To test the generalization ability of TransBind zeroshot, we evaluated it on the data of three TFs: HNF1A, ATF4, and FOXA3 that were excluded from all stages of training and validation. These factors represent distinct DNA-binding domain families: HNF1A (homeodomain), ATF4 (bZIP), and FOXA3 (forkhead).

As shown in **Table 3**, TransBind zeroshot achieves AUROC scores substantially above random (0.5) for all three unseen TFs, despite no exposure to them at all during training, indicating that it learned some general physical principles of protein-DNA interaction. In contrast, DNA sequence-only models such as TBiNet and DeepSEA completely lack this capability and would require retraining on the DNA binding data of new TFs to accommodate them. This zero-shot capability is especially valuable given that many transcription factors in both human and other species remain poorly characterized due to limited availability of ChIP-seq or other binding data. By enabling binding prediction for previously uncharacterized TFs based solely on the data of some characterized TFs, TransBind zeroshot offers a powerful tool for accelerating regulatory and functional genomics research.

**Table 3.** Zero-shot performance (AUROC) on three new transcription factors not seen during training.

| Transcription Factor | AUROC |
| --- | --- |
| HNF1A | 0.628 |
| ATF4 | 0.575 |
| FOXA3 | 0.602 |

### An Ablation Study of TransBind

To assess the impact of TF features on TransBind’s performance, we conducted an ablation study comparing two configurations under identical training conditions: the baseline TransBind, which relies solely on DNA sequence features processed through Conv1D, BiLSTM, and Transformer layers in Module 1, and the Protein-Aware TransBind, which incorporates additional TF protein features via a cross-attention mechanism. Both models were trained on the same ChIP-seq training dataset for 60 epochs, using identical hyperparameters (learning rate = 0.000328, dropout = 0.088, weight decay = 0.028, batch size = 256) with early stopping based on validation AUPR. The trained models were blindly tested on the test data. As shown in **Table 4**, integrating the TF features with DNA features yielded consistent improvements across both evaluation metrics, with AUROC increasing from 0.9493 to 0.9504 and AUPR from 0.3685 to 0.3710. The average AUPR gain, though modest, is particularly relevant for this highly imbalanced prediction task—where binding sites typically comprise a tiny portion of genomic regions. The results show that adding TF features can improve the prediction accuracy.

**Table 4.** The performance of the baseline TransBind with Protein-Aware TransBind. Values represent mean performance on the test dataset.

| Model Configuration | AUROC | AUPR |
| --- | --- | --- |
| Baseline TransBind | 0.9493 | 0.3685 |
| **Protein-Aware TransBind** | **0.9504** | **0.3710** |

Moreover, as shown in TransBind zeroshot, including TF features is required to build TF-DNA binding prediction models that can generalize to new TFs that do not exist in the training data.

## Discussion

This study addresses a key limitation in TF-DNA binding prediction: the information of TFs is not used as input to directly account for their sequence and structural properties that underlies DNA recognition. By integrating TF embeddings generated by protein language models with DNA sequence features through the cross-attention mechanism, TransBind enables TF-specific scanning of genomic sequences. This protein-aware framework delivers consistent and substantial gains in prediction accuracy over state-of-the-art deep learning methods. Significantly, it performs better in 97.1% of the 690 TF–cell type combinations tested than the second most accuracy method - TBiNet. Beyond accuracy, TransBind recovers 160 known motifs of diverse TF families, demonstrating the biological interpretability of the model and the ability to learn regulatory patterns from scratch.

Moreover, due to the use of protein embeddings of TFs as input, TransBind zeroshot also exhibits zero-shot generalization to unseen TFs, offering predictive capability for TFs with no available binding data for training, a critical step toward the scalable annotation of many uncharacterized TFs.

These improved or new capabilities have practical implications for regulatory genomics, including prioritizing TFs for experimental validation, designing synthetic regulatory sequences, and aiding variant interpretation in phenotype-associated non-coding regulatory regions in genomes.

However, the approach still has some limitations. The performance of the zero-shot prediction (i.e., AUROC between 0.575 and 0.628) is moderate, highlighting the challenge of modeling complex protein–DNA recognition without TF-specific training examples. The relatively lower performance may be largely due to the limited amount of training data of only 161 TFs. To further improve its generalization performance, a much larger training dataset involving hundreds or more TFs may be needed. In addition, the input for the zero-shot prediction in TransBind zeroshot does not implicitly contain cell type information as the multi-label classifier TransBind does, which may also lead to lower prediction accuracy.

Indeed, the current architecture of TransBind predicts TF-DNA binding from TF and DNA information only, without considering the accessibility of chrommatin and the higher-order chromatin architecture that often influences in vivo binding patterns. Moreover, protein embeddings derived from the protein language model contain only protein sequence and indirect structural information, without directly leveraging the 3D structures of TFs. Addressing these limitations will require integrating additional sources of data such as ATAC-seq chromosome accessibility data, Hi-C chromosome conformation data, predicted structures of TFs, genome annotations, genome/histone methylation, and gene expression data. It is worth noting that dynamic chromosome accessibility and gene expression data can not only enable the method to predict biological context-specific TF-DNA binding but also provide implicit cell type information for predicting cell type specific TF-DNA binding. By situating TF binding prediction within a multi-modal framework leveraging multiple sources of omics and protein structure data, more generalizable and biologically grounded models of TF-DNA binding prediction can be developed.

Finally, TransBind was currently trained on the human TF-DNA binding data. However, we image it can be trained on the TF-DNA binding data of multiple species because it can learn the general biophysical interactions between TFs and DNAs that are assumed to be universal across species. In the future, we plan to curate a large dataset consisting of TF-DNA binding data of many species and train and test TransBind on it. We will test if TransBind can generalize to TFs in new species that are not used in training at all.

## Acknowledgments

This work is supported in part by funds from the National Science Foundation (NSF: # CCF2343612 and # DBI2308699).

