## Supplementatry Note for "Integrating Protein and DNA Embeddings for Improving Genome-Wide Transcription Factor Binding Site Prediction"

### Supplementary Note S1

Shreya Basnet[1], Jianlin Cheng[1]*
[1]Electrical Engineering & Computer Science, University of Missouri, Columbia, MO 65211, USA

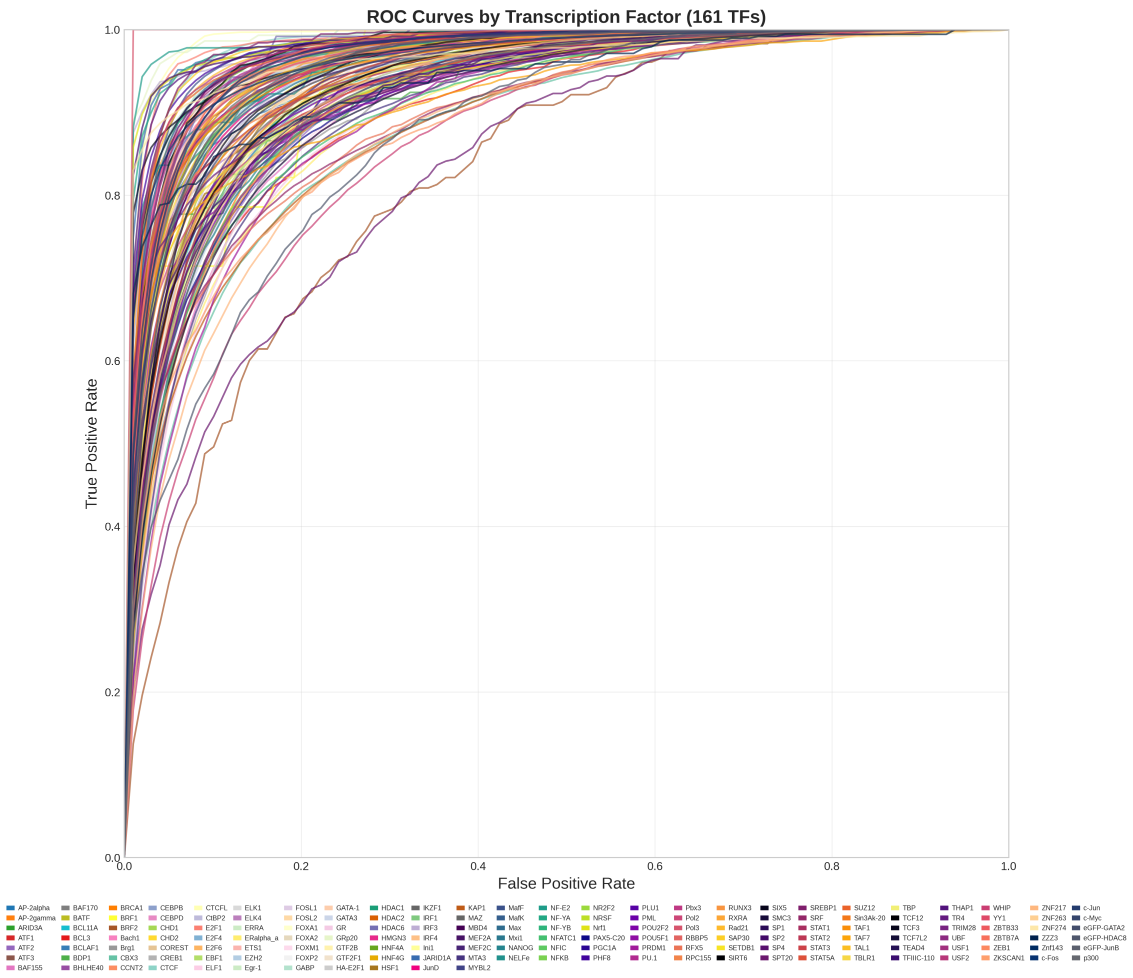


**Fig. S1.** ROC Curves for Transcription Factor Classification Performance

#### Hyperparameter Optimization

To optimize the model's performance, we conducted extensive hyperparameter tuning using the `Optuna` framework (see Supplementary Information  for details). The optimization employed the Tree-structured Parzen Estimator (TPE) sampling strategy, coupled with the `MedianPruner` for efficient early stopping of underperforming trials. Our search targeted key architectural and training parameters, including attention head counts, learning rates, regularization strengths, and network capacity configurations.

##### Search Space

- **Learning rate:** Log-uniform $[5 \times 10^{-5}, 5 \times 10^{-3}]$
- **Dropout probability:** Uniform $[0.05, 0.4]$
- **Weight decay:** Log-uniform $[1 \times 10^{-4}, 1 \times 10^{-1}]$
- **CNN output channels:** 256, 320, 384, 512
- **CNN kernel size:** 19, 26, 35, 45

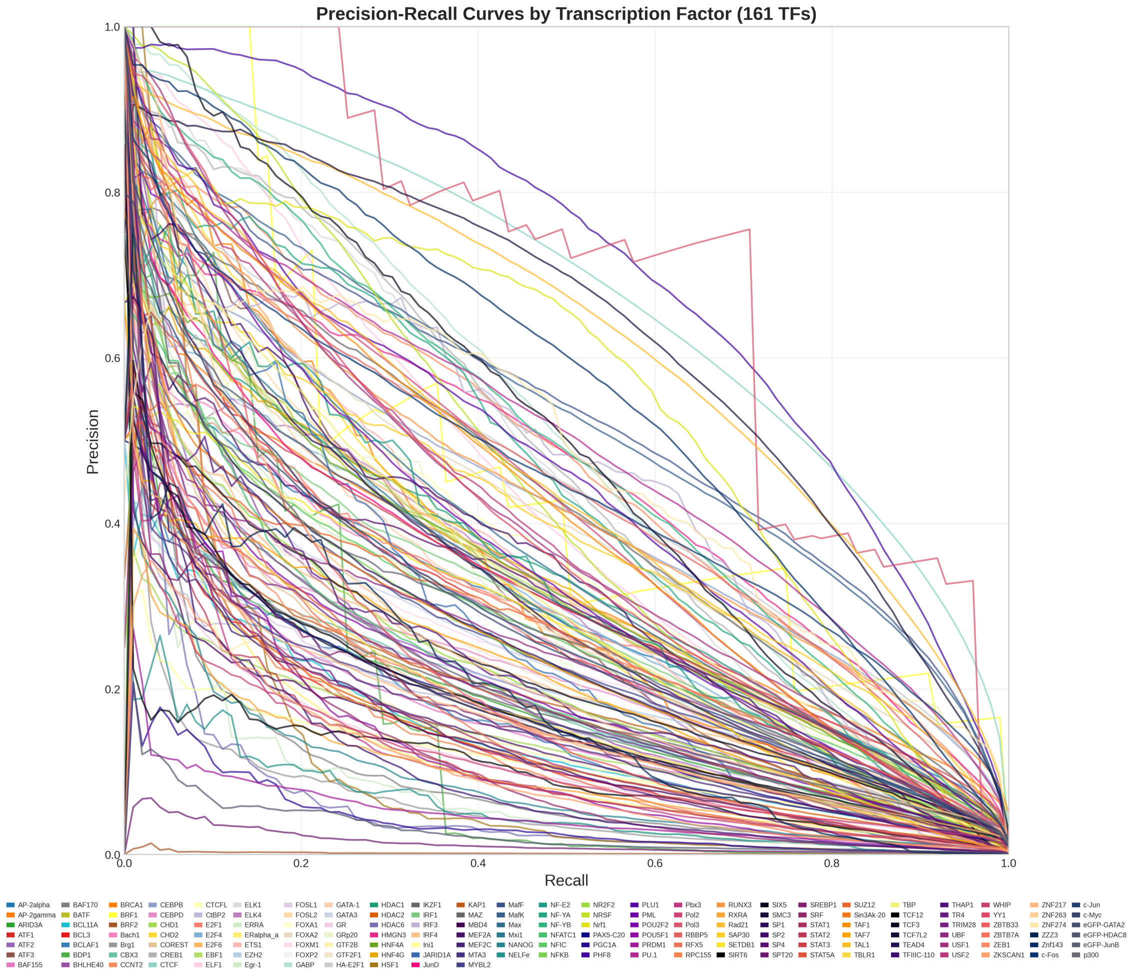


**Fig. S2.** Precision-Recall Curves for Transcription Factor Classification Performance

- **LSTM hidden size:** 128, 160, 192, 256
- **LSTM layers:** 1–3
- **Attention heads:** 4, 8, 16
- **Feedforward dim:** 512, 1024, 1280, 2048
- **FC1 size:** 512, 695, 1024, 1536

#### Training and Evaluation

Training used a fixed maximum number of epochs per configuration. TPE sampling and `MedianPruner` stopped low-performing trials early based on validation metrics. AUPR and AUROC on validation were the main optimization metrics.

#### Optimal Configuration

- Learning rate: $3.28 \times 10^{-4}$
- Weight decay: 0.028
- Dropout: 0.088
- CNN output channels: 320
- CNN kernel size: 26
- LSTM hidden size: 160
- LSTM layers: 2
- Attention heads: 16
- Feedforward dim: 1024
- FC1 size: 1024
